# The PCR Simulator: an on-line application for teaching Design of Experiments and the polymerase chain reaction

**DOI:** 10.1101/415042

**Authors:** Harold Fellermann, Ben Shirt-Ediss, Jerzy W. Koryza, Matthew Linsley, Dennis W. Lendrem, Thomas P. Howard

**Affiliations:** School of Computing, Faculty of Science, Agriculture and Engineering, Newcastle University, Newcastle-Upon-Tyne, NE1 7RU, United Kingdom; School of Mathematics, Statistics and Physics, Faculty of Science, Agriculture and Engineering, Newcastle University, Newcastle-Upon-Tyne, NE1 7RU, United Kingdom; Institute for Cellular Medicine and NIHR Biomedical Research Centre, Newcastle University, Newcastle-Upon-Tyne, NE2 4HH, United Kingdom; School of Natural and Environmental Sciences, Faculty of Science, Agriculture and Engineering, Newcastle University, Newcastle-Upon-Tyne, NE1 7RU, United Kingdom

**Keywords:** Virtual Experiments, Design of Experiments, Teaching Tools, Polymerase Chain Reaction, Web Service

## Abstract

Our PCR Simulator is a web-based application designed to introduce concepts of multi-factorial experimental design and support teaching of the polymerase chain reaction. Learners select experimental settings and receive results of their simulated reactions quickly, allowing rapid iteration between data generation and analysis. This enables the student to perform complex iterative experimental design strategies within a short teaching session. Here we provide a short overview of the user interface and underpinning model, and describe our experience using this tool in a teaching environment.

## Introduction

Over the past decades, computer simulation has become ever more integrated at all levels and in all areas of education (1). Interactive simulations can enhance the cognitive gains of learners compared with traditional learning techniques (2). Simulations are also beneficial in training for experimental procedures where long run times or high costs necessitate prescriptive practical sessions or forbid practical teaching altogether. Furthermore, experimental design and statistical analysis are highly connected activities whose teaching is rarely integrated; statistical training within the Biosciences traditionally focuses on data analysis, not experimental planning. Experimental simulations are therefore highly valuable tools to simultaneously introduce the concepts of statistical Design of Experiments (DoE) while exploring the impact that experimental design has on analytical outcome (3–5).

Our PCR Simulator is an on-line application modelled on laboratory protocols for performing the polymerase chain reaction (PCR), a ubiquitous but complex protocol for DNA amplification. While tools exist for the design of PCR primers, few tools simulate the full PCR DNA amplification cycle over time. An exception is ‘*Virtual PCR*’, a web-based service which simulates PCR following user specifications. It returns a simulated gel electrophoresis image (6). Our software is similar but offers enhancements for teaching group sessions in which learners employ the simulator to gain experience of experimental design principles. Students may select settings for 12 experimental factors (seven for the temperature cycle, four reagent volumes and one categorical choice of polymerase) with the objective of maximizing yield and purity of the amplified DNA. Here we present an overview of the implementation and interface of the simulator, modeling PCR cycles, and its use in teaching DoE.

## Methodology

### Implementation

The PCR Simulator is an on-line application implemented in python. It uses the Django framework (7) for web application architecture, scipy for numerical integration and Matplotlib and Pillow for the rendering of results. After delivering its user interface to a connected client, the server listens to incoming simulation requests, performs a virtual experiment, and returns simulation results back to the client, which is responsible for keeping track of all states. Since no user state is kept on the server, registration and login are not necessary. To provide a fluent user experience, communication between HTTP clients and the server uses AJAX requests with JSON responses. The server records requested factors and results for all incoming requests. This information is used to create a leader board where users are ranked for each result variable over a gliding window of one hour. The simulator is available at http://virtual-pcr.ico2s.org/.

### Modeling amplification of DNA by PCR

Simulating PCR experiments is performed by numerically integrating a differential equation model of the reaction, subject to the selected temperature cycle and initial reagent concentrations. The concentrations of the latter are subjected to pipetting ‘noise’ using random values within acceptable operating boundaries of commonly used pipettes. The model captures melting and annealing of all DNA species, binding and unbinding of the polymerase, primer extension and polymerase degradation. Rates of all reactions are dependent on the temperature. The model qualitatively captures parameter responses of PCR for reasonable as well as unreasonable settings while producing numerical results within seconds. For that reason, the model currently does not feature important phenomena present in PCR such as primer mis-binding and dimerization. Details of the PCR model are subject to a future publication.

## Results

### User interface

The user interface of the simulator is a single web page that combines an input form for experimental settings and a lab book that collates data from all experiments with a simulated electrophoresis gel image and a parameter table from which the user can return to any previously used settings (Fig. 1). A summary of all the user’s experiments (factors and responses) can be downloaded in CSV format for off-line analysis and experimental planning. When the simulator is in qPCR mode, the user may toggle between gel images and DNA concentration time series. The rank of the user’s yield within their cohort is displayed along with cumulative *virtual* time taken to run their experiments. The user can opt to add a username to be shown on the leader board or to appear as "anonymous" if no username is provided. This allows users to opt in or out of the competitive aspect of the tool.

**Fig. 1.**
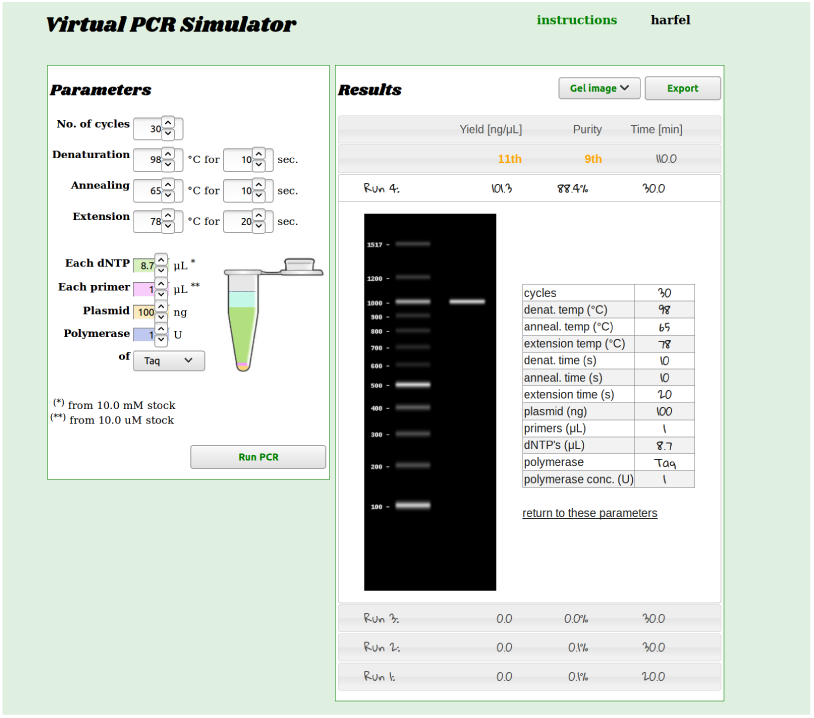
User interface. The interface comprises an input form, for experimental settings with a virtual lab book that collates results from all experiments and displays a simulated gel electrophoresis image and parameter table. The lab book also displays the rank of the user’s performance within their cohort and allows the user to export experimental settings and results in CSV format.

### Use in teaching

Over three years we have employed three iterations of the simulator to teach DoE to cohorts of approximately 20 participants from academic and industrial backgrounds with a range of laboratory and statistical experience. In-class simulator sessions break up lectures, seminars and workshop activities and each session lasts 15-30 minutes. Participants apply learning from various sessions in their interactions with the simulator, starting with free exploration of parameter space moving towards complex screening experiments. Switching from PCR to qPCR mode further increases understanding of the value of a systematic search strategy, whilst the competitive leader-board component enhances engagement. The server log can be automatically interrogated to provide information upon which an instructor might draw during a training session, or during post-teaching analysis. We have used it to assess user engagement with the task objective and to demonstrate how traditional approaches to experimentation can result in narrow search strategies (Fig. 2).

**Fig. 2.**
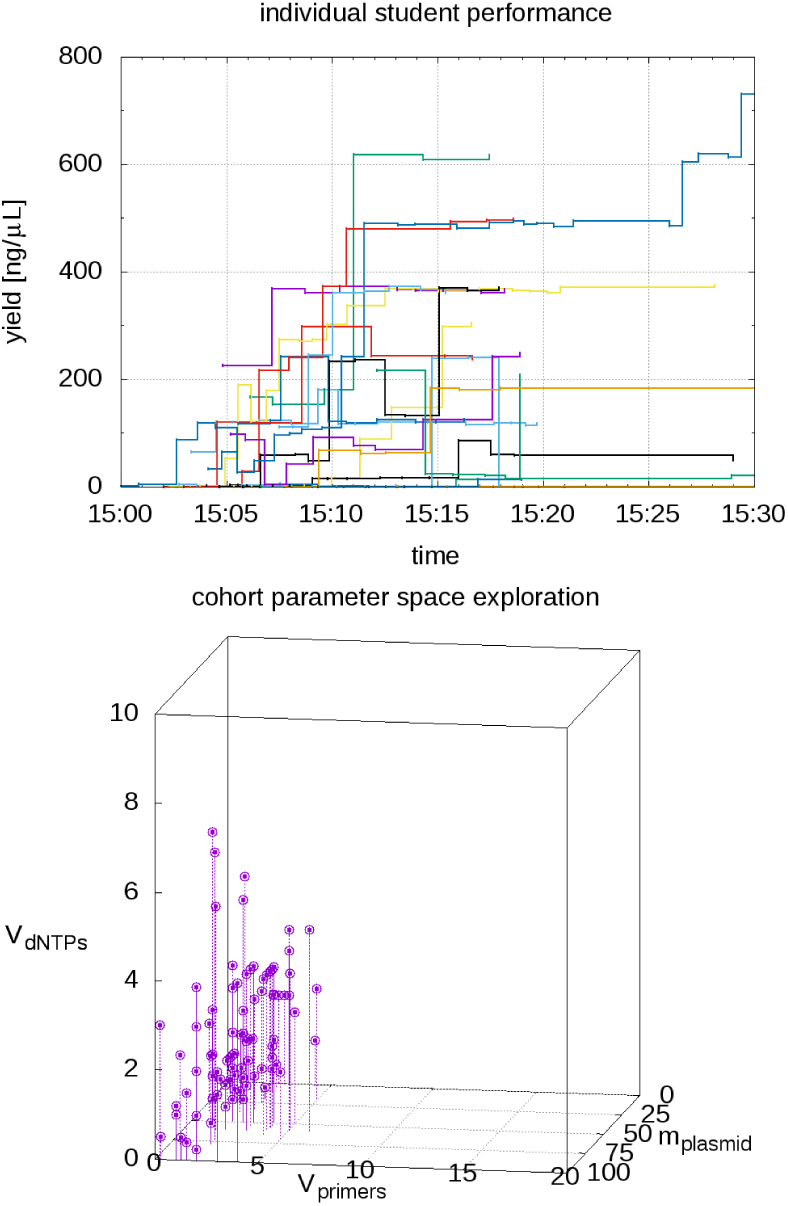
Informed teaching. The teaching interface allows the instructor to draw out information concerning student performance. This includes information on the yields and engagement of individual users (top) and how widely or effectively the cohort is exploring potential experimental options (bottom).

## Conclusion

The PCR Simulator provides a virtual experience of a complex laboratory protocol with the simulated reactions qualitatively matching real experiments. The server logs inform teacher engagement with participants and reinforce learning objectives, as well as driving a leader board to encourage engagement. Importantly, the simulator allows users to quickly generate data for their own analysis within the teaching environment allowing combined teaching of experimental design and experimental analysis

## Author contributions

HF, BSE and JWK wrote the software and mathematical model. TPH advised on principles and applications of PCR. HF, ML, DL and TPH tested and iterated the simulator in a teaching environment. All authors contributed to writing the manuscript.

## ACKNOWLEDGEMENTS

We thank Pawel Widera for his continued support as system administrator of the PCR Simulator server and a BBSRC STARS award (BB/P004342/1) for providing financial support to the workshop. BSE is supported by EPSRC (EP/N031962/1). Manuscript prepared using a modified version the HenriquesLab template available via www.overleaf.com.

